# Network-Based Biomarkers Enable Cross-Disease Biomarker Discovery

**DOI:** 10.1101/289934

**Authors:** Syed Haider, Cindy Q. Yao, Vicky S. Sabine, Michal Grzadkowski, Vincent Stimper, Maud H.W. Starmans, Jianxin Wang, Francis Nguyen, Nathalie C. Moon, Xihui Lin, Camilla Drake, Cheryl A. Crozier, Cassandra L. Brookes, Cornelis J.H. van de Velde, Annette Hasenburg, Dirk G. Kieback, Christos J. Markopoulos, Luc Y. Dirix, Caroline Seynaeve, Daniel W. Rea, Arek Kasprzyk, Philippe Lambin, Pietro Lio, John M.S. Bartlett, Paul C. Boutros

## Abstract

Biomarkers lie at the heart of precision medicine, biodiversity monitoring, agricultural pathogen detection, amongst others. Surprisingly, while rapid genomic profiling is becoming ubiquitous, the development of biomarkers almost always involves the application of bespoke techniques that cannot be directly applied to other datasets. There is an urgent need for a systematic methodology to create biologically-interpretable molecular models that robustly predict key phenotypes. We therefore created SIMMS: an algorithm that fragments pathways into functional modules and uses these to predict phenotypes. We applied SIMMS to multiple data-types across four diseases, and in each it reproducibly identified subtypes, made superior predictions to the best bespoke approaches, and identified known and novel signaling nodes. To demonstrate its ability on a new dataset, we measured 33 genes/nodes of the PIK3CA pathway in 1,734 FFPE breast tumours and created a four-subnetwork prediction model. This model significantly out-performed existing clinically-used molecular tests in an independent 1,742-patient validation cohort. SIMMS is generic and can work with any molecular data or biological network, and is freely available at: https://cran.r-project.org/web/packages/SIMMS.

---

Most human disease is complex, caused by interaction of genetic, epigenetic and environmental insults. A single disease phenotype can often arise in many ways, allowing a diversity of molecular underpinnings to yield a smaller number of phenotypic consequences. This molecular heterogeneity within a single disease is believed to underlie poor clinical trial results for some therapies^1^ and the modest performance of many genome-wide association studies^2-4^.

Researchers thus face two challenges. First, molecular markers are needed to personalize and optimize treatment decisions by predicting patient outcome (prognosis/residual risk) and response to therapy. Second, clinical heterogeneity in patient phenotypes needs to be molecularly rationalized to allow targeting of the mechanistic underpinnings of disease. For example, if a single pathway is dysregulated in multiple ways, drugs targeting the pathway could be applied.

Several approaches have been taken to solve these challenges. The most common has been to measure mRNA abundances as a snapshot of cellular state, and to construct a predictive model from them^5, 6^. Unfortunately these studies have been limited by noise and disease heterogeneity. Further, RNA is rarely directly functional^7^. Several groups have integrated multiple data-types using network and systems biology approaches identifying patient subtypes, with limited post-hoc clinical evaluation^8-26^. These algorithms have not yet clearly shown how the interplay between different pathways underpins disease etiology, nor generated biomarkers with systematically demonstrated reproducibility on independent patient cohorts across multiple indications^27^.

There is thus an urgent need to generate accurate and actionable biomarkers that integrate diverse molecular, functional and clinical information. We developed a subnetwork-based approach, called SIMMS, which uses arbitrary molecular data types to identify dysregulated pathways and create functional biomarkers. We validate SIMMS across 5 tumour types and 11,392 patients, using it to create biomarkers from a diverse range of molecular assays and uncovering unanticipated pan-cancer similarities.

## Results

### SIMMS prioritization of candidate prognostic subnetworks

SIMMS acts upon a collection of subnetwork modules, where each node is a molecule (*e.g*. a gene or metabolite) and each edge is an interaction (physical or functional) between those molecules. Molecular data is projected onto these subnetworks using topology measurements that represent the impact of and synergy between different molecular features. To allow modeling of biological processes with different network architectures, we devised three network topology measurements: (nodes/molecules only), E (edges/interactions only) and N+E (nodes and edges). While the N model assumes independent and additive effects of parts of a subnetwork, the E and N+E models incorporate the impact of dysregulated interactions (**Online Methods**). SIMMS fits one of these models and computes a ‘module-dysregulation score’ (MDS) for each subnetwork that measures their strength of association with a specific disease, phenotype or outcome (**Supplementary Figure 1**).

### Characteristics and benchmarking of SIMMS identified prognostic subnetworks

A key challenge faced by translational research is to extend the single gene biomarkers paradigm to clinically actionable metagenes/pathways. Thus, we tested the prognostic value of pathway-derived subnetworks using Cox modeling to quantify how effectively a subnetwork stratifies patients into groups with differential risk (**Online Methods**). SIMMS can use any network, and we chose to evaluate it using 449 gene-centric pathways from the high-quality, manually-curated NCI-Nature Pathway Interaction database^28^. These pathways comprise 500 non-overlapping subnetworks (**Supplementary Table 1**; **Supplementary Figure 2**). We then trained and tested SIMMS on a series of large and well-curated mRNA abundance datasets of primary breast (1,010 training patients; 1,098 validation patients), colon (205 training; 439 validation), lung (380 training; 369 validation) and ovarian (438 training; 566 validation) cancers (**Supplementary Tables 2-5**; **Supplementary Figure 3**).

Our analysis of prognostic subnetworks revealed several properties of tumour network biology. First, there was a global propensity for highly prognostic subnetworks to contain significantly higher number of genes and interactions for Model N and N+E (P<0.05, Wilcox rank sum test; **Supplementary Figure 4**). This association between subnetwork size (number of genes) and prognostic power was consistent in breast, NSCLC and ovarian cancers, even though different pathways were altered in each but not in colon cancers. Second, the prognostic ability of Model N was significantly superior to that of Model N+E and E; a trend which was maintained across all four cancer types (one-way ANOVA, Tukey HSD multiple comparison test; **Supplementary Figure 5)**. This suggests that mRNA abundance of functionally-related genesets alone is a strong predictor of patient outcome. We therefore focused solely on Model N moving forward, while recognizing that in other diseases different regulatory architectures may be disrupted.

Next we compared how SIMMS subnetwork scores perform against five well-known machine learning algorithms treating genes as individual features in multivariate setting (**Supplementary Results section 2**). SIMMS identified an equal or significantly greater number of prognostic subnetworks compared to models based on genes in each of these subnetworks for these methods (P<0.01, proportion test; **Figures 1a-d**).

**Figure 1.**
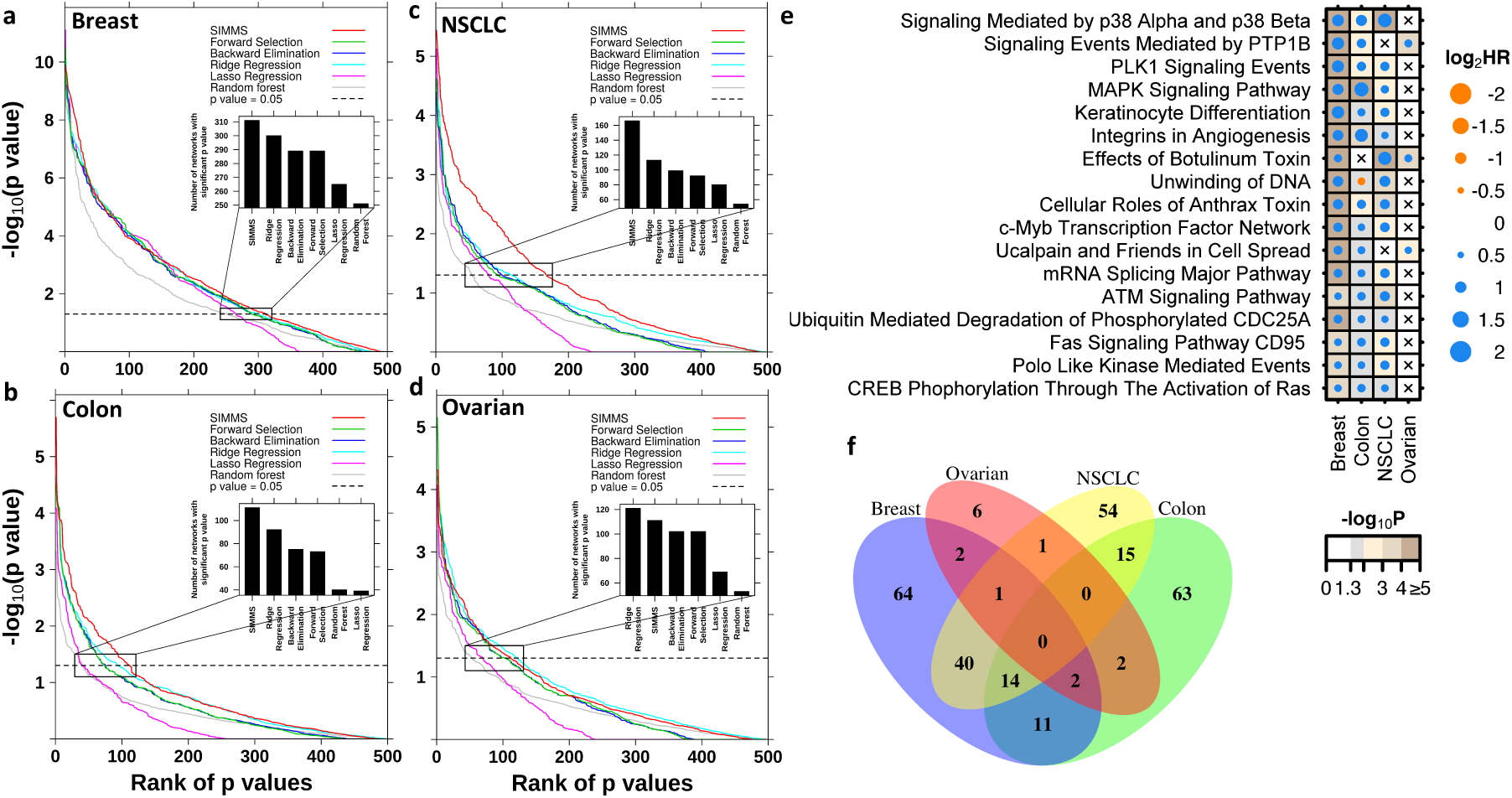
Pan-cancer prognostic subnetworks. **(a)** Comparison of prognostic assessment of subnetworks in validation sets of breast cancer using SIMMS and five machine learning algorithms. For each algorithm, p values were ranked with rank 1 assigned to the smallest P value. Number of validated subnetworks identified by each algorithm (P < 0.05, above horizontal dashed line) are shown as barplots. **(b-d)** Same as (a) but for colon, NSCLC and ovarian cancers. **(e)** Dot plot of univariate hazard ratios and P values (Wald-test) for each of the top *n* subnetworks significantly associated with patient outcome (|log_2_ HR| >1.5, P < 0.05) in at least 3/4 cancer types. A Cox proportional hazards model was fitted to dichotomized risk scores across the entire validation cohort. Crosses represent absence of a module from a particular cancer type. **(f)** Overlap of candidate subnetwork markers across breast, colon, NSCLC and ovarian cancers.

### Multi-cancer analysis reveals recurrently dysregulated subnetworks

We next quantitatively determined the similarity between different tumour types at the pathway level. Cross disease assessment of significantly prognostic subnetworks (P<0.05) revealed well-known oncogenic pathways such as Aurora Kinase A and B signaling, apoptosis, DNA repair, *RAS* signaling, telomerase regulation and *P53* activity in all four tumour types (**Supplementary Tables 6-9**). By limiting to highly prognostic subnetworks (|log_2_HR|>1.5 and P<0.05) in each tumour type, 17 recurrently prognostic subnetworks (at least three tumour types) were identified (**Figure 1e, Supplementary Figure 6**). Significant overlap between prognostic subnetworks was observed for breast, colon and NSCLC (14 subnetworks: P_overlap_=0.009, 10^5^ permutations; **Figure 1f**). These results can inform prospective clinical trials on repurposing strategies of anti-cancer drugs targeting the pathways underlying these subnetworks.

In breast cancer, subnetworks modules encompassing proliferation pathways (Mitosis, *PLK1, AURKA* and *AURKB*) were highly prognostic (**Supplementary Table 6b**). To ensure these are not driven by common proliferation genes, we tested gene overlap in these subnetworks and found them highly divergent (**Supplementary Figure 7a**). We further tested whether estimated risk-scores recapitulate proliferation accurately. We used the *MKI67* (mRNA abundance) as a surrogate for proliferation and showed strong concordance with SIMMS risk-scores (Spearman’s *ρ*=0.79-0.86, P<10^-3^; **Figure 2a**). To determine if subnetworks more accurately model patient-relevant biology, we constructed a multivariate proliferation signature using the four modules. This signature was a robust prognostic marker (**Figure 2b)** and presents an opportunity to understand the functionally-related proliferation correlates of patient outcome beyond single gene markers.

**Figure 2.**
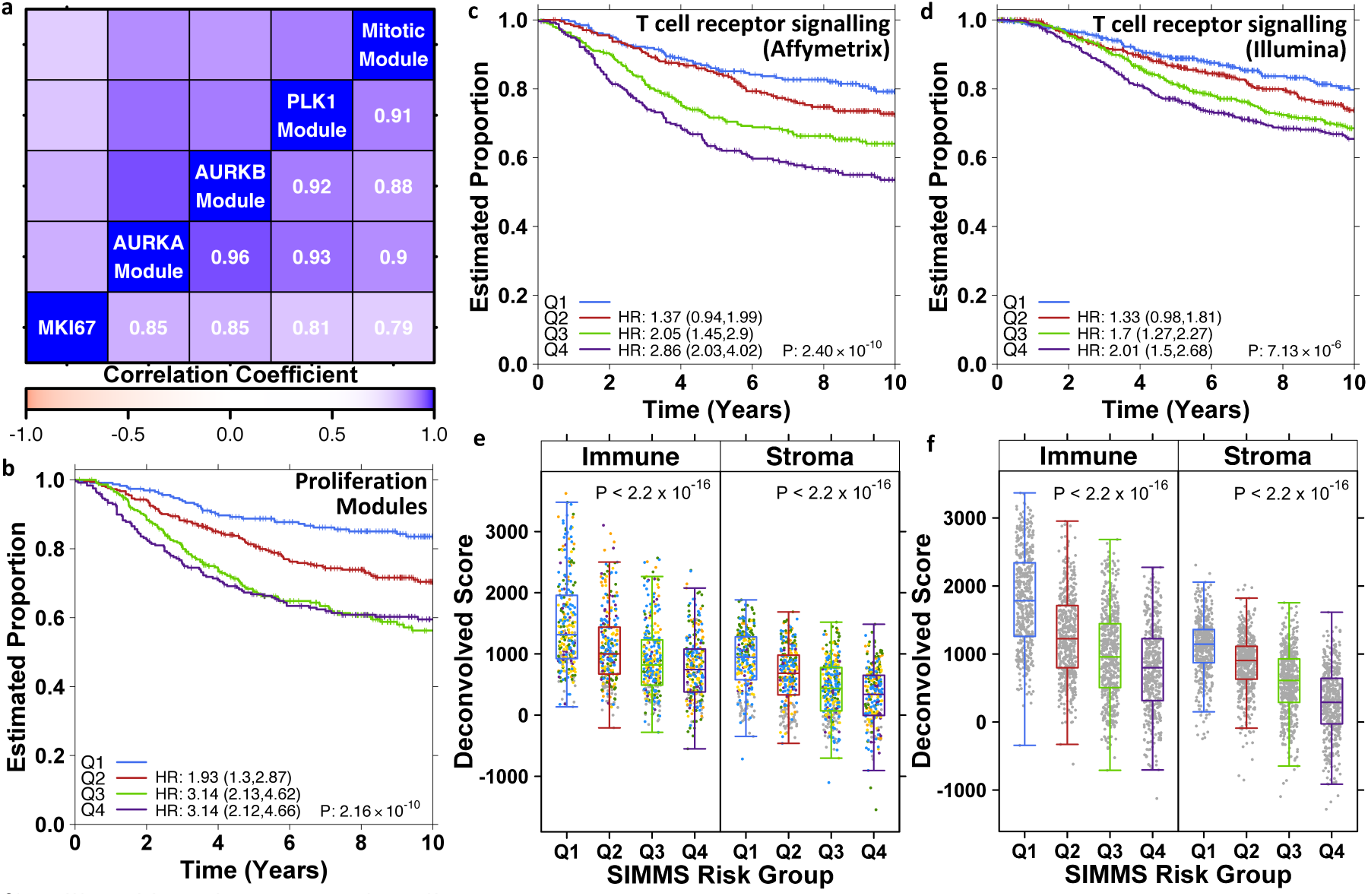
Proliferation and immuno subnetworks. **(a)** Heatmap of correlation (Spearman) and cluster analysis of patient’s risk-scores for proliferation modules in breast cancer, alongside mRNA abundance of a proliferation marker *MKI67*. Data shown is from validation cohorts. **(b)** Kaplan-Meier analysis of predicted proliferation scores (validation cohorts) using SIMMS-derived proliferation biomarker. Groups (Q1-Q4) were established using quartiles derived from the training set, and groups Q2-Q4 were compared to Q1 using Cox proportional hazards model. P value was estimated using Log-rank test. **(c)** Kaplan-Meier analysis of tumour immune microenvironment driver subnetwork (BioCarta pathway: T cell receptor signaling) in Affymetrix based validation cohorts. Quartile based risk groups (thresholds derived using training set cohorts), demonstrating linear increase in the likelihood of recurrence/event. Test statistics estimation same as in **(b). (d)** Kaplan-Meier analysis of tumour immune microenvironment driver subnetwork (BioCarta pathway: T cell receptor signaling) in Metabric breast cancer cohort (Illumina). **(e)** Assessment of computationally inferred immune system infiltration and stromal estimates against SIMMS predicted risk groups (Q1-Q4 *i.e*. low to high) in Affymetrix validation cohorts (test statistics shows P value of ANOVA). Colour of dots represent respective validation cohort (**Supplementary Table 2). (f)** Same as **(e)** in Metabric cohort (Illumina).

We next investigated prognostic subnetworks focusing on clinically actionable pathways. In breast cancer, Immune microenvironment subnetwork of *T cell receptor signaling* was significant predictor of patient outcome (HR_Q1-Q4_=2.86, 95% CI=2.03-4.02, P=1.78 × 10^-9^; **Figure 2c, Supplementary Table 6d**), in particular distant metastasis free survival where data was available (Sotiriou: HR=3.52, 95% CI=1.38-9.02, P=0.0086; Wang: HR=1.58, 95% CI=1.07-2.33, P=0.02). We further validated this subnetwork for breast cancer disease-specific survival in an independent cohort of 1,970 patients^29^ (HR_Q1-Q4_=2.01, 95% CI=1.5-2.68, P=7.13 × 10^-6^; **Figure 2d**). Hypothesizing that this subnetwork may serve as a marker of tumour immune infiltration, we confirmed association between SIMMS predicted risk groups and immune cell content^30^ (Affymetrix: Spearman’s *ρ*=-0.38, P < 2.2 × 10^-16^; Illumina: Spearman’s *ρ*=-0.48, P < 2.2 × 10^-16^) as well as stromal signal (Affymetrix: Spearman’s *ρ*=-0.43, P < 2.2 × 10^-16^; Illumina: Spearman’s *ρ*=-0.59, P < 2.2 × 10^-16^) (**Figures 2e-f**), both of which were associated with good outcome. Consistent with a recent breast cancer study^31^, naïve immune and stromal content estimates were only weakly associated with patient outcome (**Supplementary Figures 7b-e**), whilst SIMMS’s MDS of *T-cell receptor signaling* not only provides accurate identification of patients who may benefit from immunotherapy but also indicates associated molecular targets.

### Subnetwork-based biomarkers predict patient outcome

As SIMMS accurately identified individual prognostic subnetworks, we hypothesized that modeling of multiple aspects of tumour biology through these subnetwork into a single molecular biomarker could better rationalise patient heterogeneity emerging from alternative pathways of disease progression. First, to initialise the number of subnetworks, 10 million random sets of subnetworks of different sizes (1 to 250) were generated. These were tested for prognostic potential in a multivariate Cox model, thereby generating an empirical null distribution which allowed us to select the optimal number of pathways that influence survival in each disease (**Supplementary Figure 8**). Using the optimal size of subnetworks maximizing performance in the training set (n_Breast_ = 50, n_Colon_ = 75, n_NSCLC_ = 25 and n_Ovarian_ = 50), SIMMS risk-scores was estimated for top n subnetworks in each disease. These subnetworks revealed a number of highly correlated clusters of subnetworks (**Supplementary Figures 9-12)**. Next, multivariate prognostic classifiers (Cox model with *L1* regularization; 10-fold cross validation) were created for each tumour type thereby further refining the list of highly correlated subnetworks. For each tumour type, subnetwork-based classifiers encompassing multiple pathways successfully predicted patient survival in the fully-independent validation cohorts (**Figure 3, Supplementary Tables 10-13**). We verified that these results are not driven by a single cohort or patient subset, but rather reproducible across studies (**Supplementary Figures 13-16**). Similarly SIMMS generated robust biomarkers for each tumour-type using multiple feature-selection approaches: multivariate analysis using both backward and forward refinements yielded similar results (**Supplementary Figure 17**). We further challenged SIMMS’s paradigm of selection of the top n pathway-based features for multivariate modelling against biomarker constructed from all genes in multivariate setting using a Cox model with *L1* regularization. While breast and ovarian cancers yielded results similar to SIMMS; colon and NSCLC models were significantly inferior to SIMMS’s models (**Supplementary Figure 18**). To ensure SIMMS-derived prognostic markers performed comparably to existing transcriptomic prognostic tools, we compared our four SIMMS signatures to 21 independent approaches in the same test datasets. For each disease, the SIMMS signature performed as-well or better than the best published signature, each of which used a unique methodology (**Supplementary Results section 3, Supplementary Table 14**). Therefore SIMMS provided a consistent and unified approach to generating highly accurate biomarkers.

**Figure 3.**
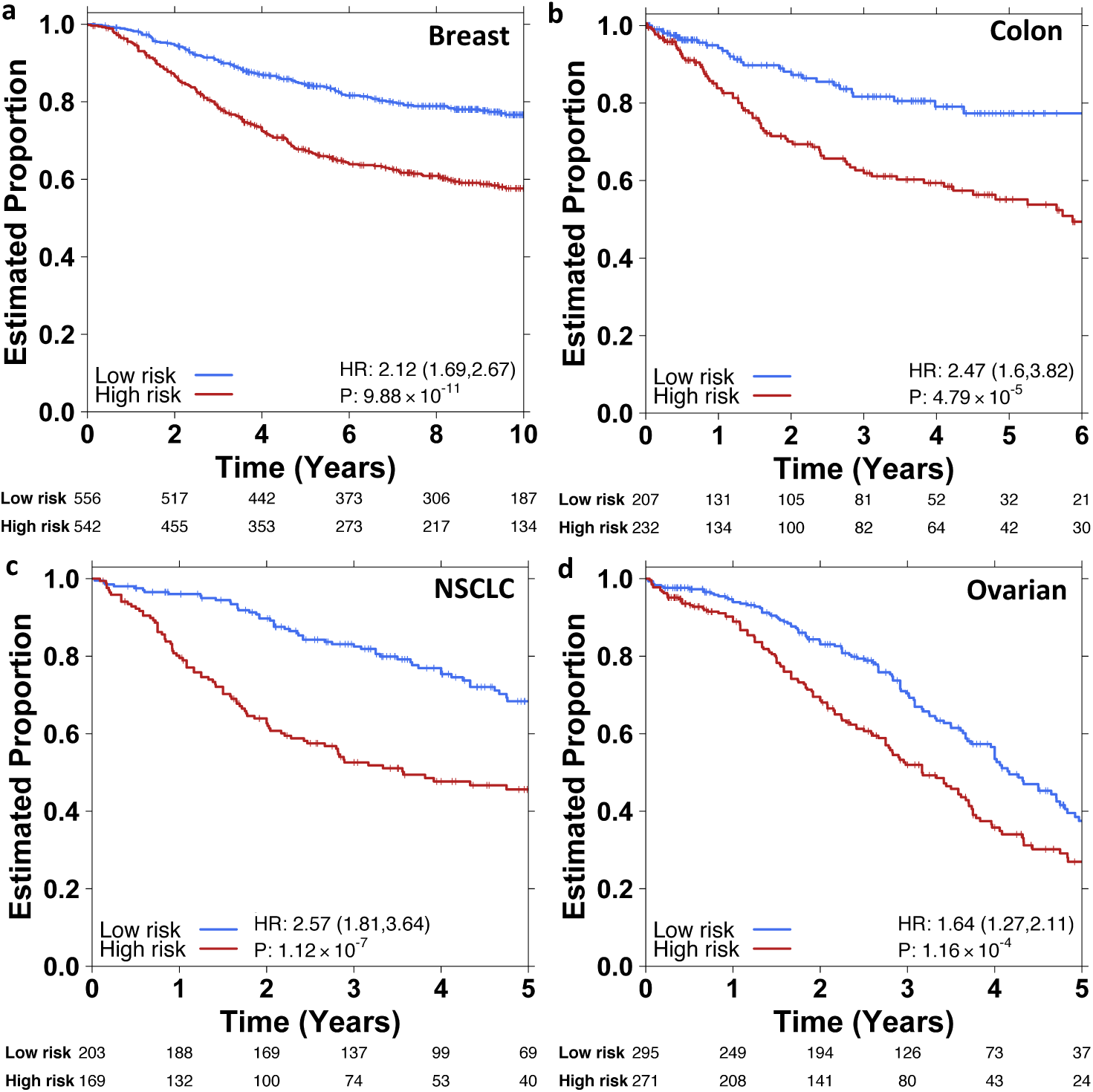
SIMMS biomarkers for multiple tumour types. **(a-d)** Kaplan-Meier survival plots using Model N over the entire validation cohort with subnetwork selection conducted through Cox model based Generalised Linear Models (*L1*-penalty) on the training cohort. Final model resulted in 23/50, 5/75, 23/25 and 23/50 subnetworks for breast, colon, NSCLC and ovarian cancers respectively (**Supplementary Tables 10-13**). P values were estimated using Wald-test.

Focusing on breast cancer as a disease with well-defined molecular subtypes, we tested SIMMS on Metabric breast cancer cohort (n=1,970)^29^. Our prognostic classifier revealed two primary patient clusters with distinct pathway activities. These clusters were highly correlated with the PAM50^32^ intrinsic subtypes of breast cancer (*f*-measure=0.81; **Figure 4a**). Since breast cancer is a heterogeneous disease with distinct molecular and clinical characteristics^32^, we asked whether SIMMS could identify subtype-specific prognostic markers. To evaluate this, we classified patients into PAM50, ER+ and ER-subtypes and created SIMMS classifiers for each subtype. SIMMS classifiers were able to identify subgroups of patients at a significantly higher risk of relapse (P<0.05) in each of the Luminal-A, Normal-like and ER+ subtypes (**Figure 4b**). Importantly, these subgroups of patients present differential pathway activity (as quantified by SIMMS), and hence may benefit from aggressive/alternative treatments targetting these pathways. We further validated the efficacy of SIMMS when trained and tested for reproducibulity across different genomic platforms (Affymetrix and Illumina; P<10^-5^; **Figure 4b AFFY/ILMN, ILMN/ILMN, ILMN/AFFY; Supplementary Figure 19**). Taken together these results demonstrate that pathway-driven subnetwork modelling can flexibly integrate diverse assays emerging from multiple platforms.

**Figure 4.**
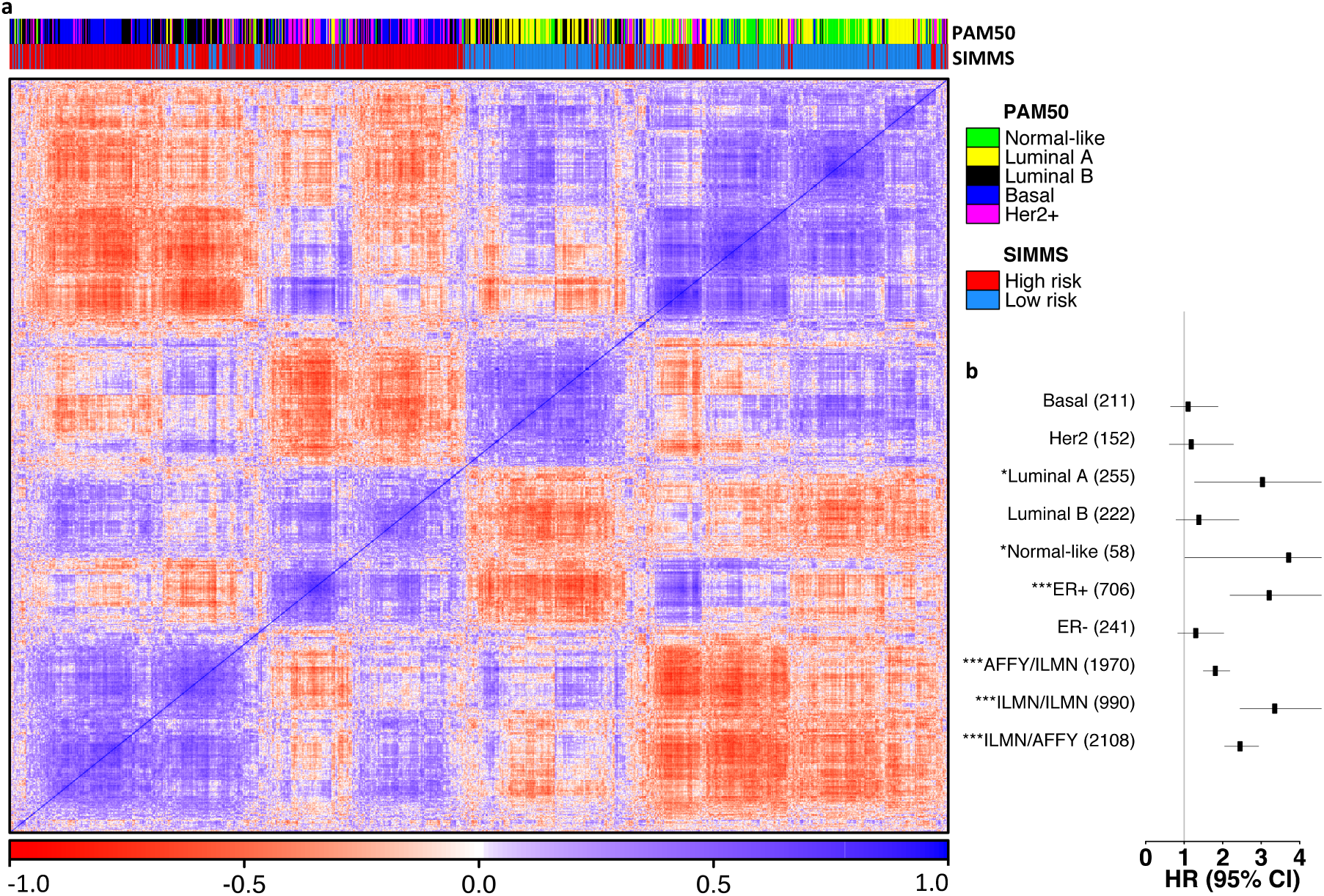
Clinical analysis of breast cancer biomarkers. **(a)** Heatmap of correlation and cluster analysis of patients’ risk-scores of top n_Breast=_50 subnetworks in the Metabric validation cohort. The covariates show concordance (estimated using *f*-measure) between PAM50-based molecular subtypes and SIMMS predicted risk group. **(b)** Forest plot showing HR and 95% CI (multivariate Cox proportional hazards model) of the breast cancer subtype-specific markers as well as cross-platform validation. Datasets originating from Illumina (ILMN) and Affymetrix (AFFY) were used in turn for cross platform training and validation purposes. Due to limited availability of clinical annotations on Affymetrix based cohorts, only the Illumina dataset (Metabric) was used for subtype-specific models. For these, the Metabric-published training and validation cohorts were maintained for training and validation purposes. Numbers in parenthesis indicate the size of the validation cohort. Asterisks represent statistical significance of differential outcome between the predicted low- and high-risk groups (* P<0.05, ** P<0.01, *** P<0.001, Wald-test).

### A PIK3CA signaling residual risk predictor in early breast cancer

While the public data used to evaluate SIMMS is valuable, it does not closely represent that used in clinical settings. To better represent this scenario, we focused on the PI3K-signaling pathway, which is frequently mutated in breast cancer and is the subject of several targeted therapies. We evaluated 1,734 samples from the Phase III TEAM clinical trial and measured mRNA abundance of 33 PI3K signaling genes in clinically-relevant FFPE samples. All samples were ER positive “luminal” breast cancers from the TEAM pathology study^33^ (**Supplementary Table 15, Supplementary Results section 4**). We hypothesized that inclusion of key signaling nodes from driver molecular pathways in residual risk signatures would both improve risk stratification and identify candidate theranostic targets for the next generation of clinical trials. Univariate prognostic assessment of 33 genes revealed significant association between seven genes and distant metastasis (Wald P_adjusted_<0.05; **Supplementary Table 16**). Survival analysis of clinical covariates indicated tumour grade, N-stage, T-stage and HER2 IHC as predictors of distant metastasis (**Supplementary Table 17**). Next, we aggregated 33 PI3K signaling genes into 8 functional modules representing different nodes of the pathway (**Supplementary Figure 20, Supplementary Table 18**), and applied SIMMS to train a residual risk model. The SIMMS-derived model comprised of four modules and two clinical covariates (**Supplementary Table 19**).

To validate this model, we profiled using the same technologies and gene set a fully-independent set of 1,742 patients from the same clinical trial (**Supplementary Table 20**). This scenario closely replicates actual clinical application of the signature. The SIMMS signature was a robust predictor of distant metastasis in the validation cohort (**Figure 5a**; Q4 *vs*. Q1 HR=9.68, 95%CI: 5.91-15.84; P=2.22×10^-40^). It was also effective when simply median-dichotomizing predicted risk scores into low- and high-risk groups (**Supplementary Figure 21a**). Risk scores from this signature were directly correlated with the likelihood of distant recurrence at five years, with a higher risk score associated with a higher likelihood of metastasis (**Figure 5b**). The signature was independent of PIK3CA point mutations, with no change in survival curves between low and high-risk groups with *vs*. without PIK3CA mutations (p_low+/-_=0.22, p_high+/-_=0.81; **Supplementary Figure 21b**). The signature remained an independent prognostic indicator following adjustment for chemotherapy (Q4 *vs*. Q1 HR=9.88, 95%CI: 6.01-16.27; P=2.22×10^-40^). To further verify this, predicted risk groups (Q1-Q4) in the validation cohort were divided into chemotherapy negative and positive arms with further stratification by nodal status. Risk predictions was similar for node-negative/chemotherapy-negative patients (Q4 *vs*. Q1 HR=7.69, 95%CI: 3.19-18.58; P=7.71×10^-6^; **Supplementary Figure 21c**) as for node-positive/chemotherapy-negative patients (Q4 *vs*. Q1 HR=8.76 95%CI: 3.78-20.29; P=4.24×10^-19^; **Supplementary Figure 21d**).

**Figure 5.**
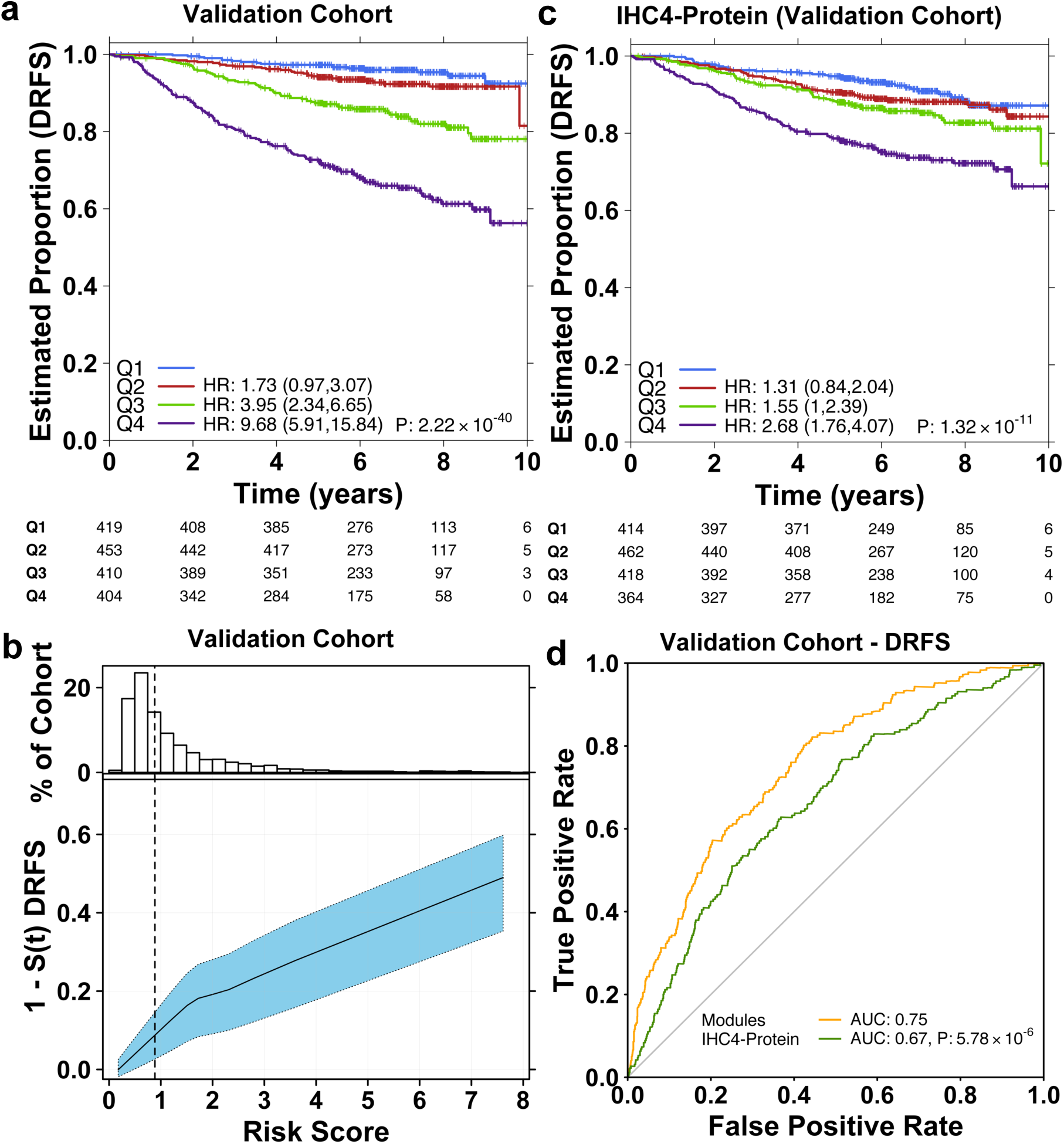
PI3KCA signaling predictor of breast cancer recurrence. **(a)** Independent validation of prognostic model trained on SIMMS’s risk-scores and clinical covariates (N and tumor size). Risk score estimates were grouped into quartiles derived from the TEAM training cohort; each group was compared against Q1. Hazard ratios were estimated using Cox proportional hazards model and significance of survival difference was estimated using the log-rank test. **(b)** Distribution of patient risk scores in the TEAM Validation cohort (top panel). Bottom panel shows the predicted 5 year recurrence probabilities (solid line) and 95% CI (dashed lines) as a function of patient risk score. Vertical dashed black line indicates training set median risk score. **(c)** Risk prediction by the IHC4 protein model in the validation cohort. Quartiles were defined in the training cohort and applied to the validation cohort. Quartiles Q2-Q4 were compared against Q1, with adjustment for age, Nodal status, tumor size and grade using Cox proportional hazards modelling and the log-rank test. **(d)** Comparison of SIMMS’s modules model and IHC4-protein model using area under the *receiver operating characteristic* (AUC) curve as performance indicator.

### PIK3CA signaling modules outperform existing markers

To benchmark SIMMS’s PI3K modules signature against current clinically-validated approaches, we compared its performance to a clinically-used protein-based residual risk predictor, IHC4^34^. IHC4 was assessed using quantitative IHC measurements of ER, PgR, Ki67 and HER2^35^ with adjustment for age, nodal status, grade and size in both the training and validation (P=1.32×10^-11^; **Figure 5c**) cohorts. To compare the two predictors, we used the area under the *receiver operating characteristic* curve as a performance indicator. The PI3K modules model (AUC=0.75) was significantly superior to the IHC-protein model (AUC=0.67; P=5.78×10^-6^; **Figure 5d**). The PIK3CA predictor correctly identified 78.7% (NPV=0.93, PPV=0.27) of patients with disease relapse compared to 63.0% (NPV=0.88, PPV=0.22) by IHC4 in the validation cohort. Overall, it improved patient classification relative to IHC4 for 18% of patients (Net reclassification index = 0.18, 95% CI = 0.11-0.25, P < 2.2×10^-16^).

### General multimodal biomarkers

Since oncogenic insults manifest across all molecular species (*e.g*. DNA, mRNA, *etc*.), there is a need to simultaneously integrate these into a unified predictive models. We used four TCGA datasets (breast (BRCA)^36^, colorectal (COADREAD)^8^, kidney (KIRC)^33^, ovarian (OV)^37^) along with the Metabric^29^ breast cancer cohort, each of which included matched mRNA, CNA and clinical data, along with MEMo pathway modules published in TCGA studies (**Supplementary Table 21**). SIMMS risk-scores were estimated for each of the mRNA and CNA profiles with subnetwork weights of constituent genes calculated independently. The sum of mRNA and CNA MDS yielded a multi-modal pathway activation estimate per patient (**Supplementary Results section 5**). Multi-modal markers of kidney (5/7) and breast (19/23) cancers were reproducibly superior (Fisher’s combined probability test) to both mRNA- and CNA-alone (**Supplementary Figure 22a:** dark brown dots against red and blue covariates, **Supplementary Figures 22b-c**). For ovarian cancer, multi-modal markers improved upon CNA models in 2/3 subnetworks (**Supplementary Figure 22a:** M2 and M3 against purple covariate, **Supplementary Figures 22b-c**) even though no individual data type was prognostic in all subnetworks. These results demonstrate the potential of network models to create integrated multi-modal biomarkers.

## Discussion

Patients with complex human diseases present highly heterogeneous molecular profiles, ranging from a few aberrant genes to a set of dysregulated pathways. Because many different molecular aberrations can give rise to a single clinical phenotype, the importance of generating multi-modal datasets is increasingly appreciated^8, 29, 37, 38^. Indeed, a single whole-genome sequencing experiment generates information about single nucleotide variations, copy number aberrations and genomic rearrangements. SIMMS puts this molecular variability into the context of existing knowledge of biological pathways using subnetwork information. Several other groups have considered the value of network models in predicting breast cancer outcome^11, 12, 39^ and in subtyping glioblastoma^40^. However no such tools have yet been developed to be generalizable to a broad range of diseases or to arbitrary topological measures that might be used to estimate interaction weights in network-models of biology^41, 42^ or to work with physical, functional, transcriptional or metabolic networks^43, 44^. SIMMS provides this generalizability and flexibility by treating molecular profiles as generic featuers and not just genes.

Most previous biomarker studies have focused on establishing biomarkers using mRNA abundance profiles, with pathway-level analysis used *post hoc* to characterize the most interesting genes^45-47^. Our approach inverts this strategy, taking known pathways *a priori* and thus creating immediately interpretable and clinically actionable biomarkers^13^. For example, our PIK3CA risk predictor (**Figure 5a)** serves as a candidate assay for patient stratification in theranostic clinical trials. Both the IHC4 and type I receptor tyrosine kinase modules have extensive clinical and pre-clinical data validating their utility in early breast cancer^48, 49^. The documented effects of PIK3CA pathway inhibitors in advanced breast cancer, if appropriately targeted, may be translated into significant improvements in survival in early breast cancer.

Precision molecular medicine is predicated on the concept of giving each patient the right drug in the right dose at the right time. This type of personalized treatment requires the development of robust biomarkers that precisely predict clinical phenotypes. Current clinical biomarkers are typically derived from a small number of genes, and do not yet recapitulate the full complexity of disease. SIMMS takes a step towards integrating diverse cellular processes into a singular model, and is well-positioned to take into account the influx of clinical sequencing data now being generated. However, as -omic techniques evolve to rapidly analyze and quantify cellular metabolites, network models may need to change from being gene-centric to including metabolites as core nodes. Further, single-cell analysis methods may allow accurate interrogation of the interactions between different cell-types, perhaps requiring simultaneous fitting of multiple distinct, but interacting network models. The continued development of robust, general biomarker discovery algorithms is thus required to generate the accurate and reproducible biomarkers needed for transforming medical care.

## Online Methods

### Pathways data-preprocessing

The pathway dataset was downloaded from the NCI-Nature Pathway Interaction database^28^ in PID-XML format (**Supplementary Table 1**). The XML dataset was parsed to extract protein-protein interactions from all the pathways using custom Perl (v5.8.8) scripts (**Supplementary File 1**). The protein identifiers extracted from the XML dataset were further mapped to Entrez gene identifiers using Ensembl BioMart (version 62). Whereever annotations referred to a class of proteins, all members of the class were included in the pathway, in some case using additional annotations from Reactome and Uniprot databases. The protein-protein interactions, once mapped to the Entrez gene identifiers, were grouped under respective pathways for subsequent processing. The initial dataset contained 1,159 variable size subnetworks (**Supplementary Figure 2a-b**). In order to identify redundant subnetworks, we tested the overlap between all pairs of subnetworks. When a pair of subnetworks had a two-way overlap above 80% (if two modules shared over 80% their network components; nodes and edges), we eliminated the smaller module. Additionally, all subnetworks modules containing less than 3 edges were excluded. In total, these criteria removed 659 subnetworks, resulting in 500 subnetworks.

### mRNA abundance data pre-processing

All pre-processing was performed in R statistical environment (v2.13.0). Raw datasets from breast, colon, NSCLC and ovarian cancer studies (**Supplementary Tables 2-5**) were normalized using RMA algorithm^50^ (R package: affy v1.28.0) except for two colon cancer datasets (TCGA and Loboda dataset) which were used in their original pre-normalized and log-transformed format. ProbeSet annotation to Entrez IDs was done using custom CDFs^51^ (R packages: hgu133ahsentrezgcdf v12.1.0, hgu133bhsentrezgcdf v12.1.0, hgu133plus2hsentrezgcdf v12.1.0, hthgu133ahsentrezgcdf v12.1.0, hgu95av2hsentrezgcdf v12.1.0 for breast cancer datasets. hgu133ahsentrezgcdf v14.0.0, hgu133bhsentrezgcdf v14.0.0, hgu133plus2hsentrezgcdf v14.0.0, hthgu133ahsentrezgcdf v14.0.0, hgu95av2hsentrezgcdf v14.0.0 and hu6800hsentrezgcdf v14.0.0 for the respective colon, NSCLC and ovarian cancer datasets). The Metabric breast cancer dataset was preprocessed, summarized and quantile-normalized from the raw expression files generated by Illumina BeadStudio. (R packages: beadarray v2.4.2 and illuminaHuman v3.db_1.12.2). Raw Metabric files were downloaded from European genome-phenome archive (EGA) (Study ID: EGAS00000000083). Data files of one Metabric sample were not available at the time of our analysis, and were therefore excluded. All datasets were normalized independently. TCGA breast (BRCA), colon (COADREAD), kidney (KIRC) and ovarian (OV) cancer datasets were downloaded from http://gdac.broadinstitute.org/ (Illumina HiSeq rnaseqv2 level 3 RSEM; release 2014-01-15). The choice of training and validation sets was driven by maintaining homogeneity in size and platforms, and was further addressed through 10-fold cross validation as well as permutation analyses. Raw mRNA abundance NanoString counts data were pre-processed using the R package NanoStringNorm^52^ (v1.1.16; **Supplementary Results section 4**). A range of pre-processing schemes was assessed to optimize normalization parameters (**Supplementary Results section 4**).

### TEAM study population

The TEAM trial is a multinational, randomized, open-label, phase III trial in which postmenopausal women with hormone receptor-positive luminal^53^ early breast cancer were randomly assigned to receive exemestane (25 mg once daily), or tamoxifen (20 mg once daily) for the first 2.5-3 years followed by exemestane (total of 5 years treatment). This study complied with the Declaration of Helsinki, individual ethics committee guidelines, and the International Conference on Harmonization and Good Clinical Practice guidelines; all patients provided informed consent. Distant metastasis free survival (DRFS) was defined as time from randomization to distant relapse or death from breast cancer^53^.

The TEAM trial included a well-powered pathology research study of over 4,500 patients from five countries (**Supplementary Table 15**). Power analysis was performed to confirm the study size had 98.57% and 98.82% power to detect a HR of at least 2 in the training and validation cohorts respectively (**Supplementary Results section 4**) analyses and statistical methods followed REMARK guidelines^54^. After mRNA extraction and NanoString analysis, 3,476 samples were available. Patients were randomly assigned to either a training cohort (n=1,734) or the validation cohort (n=1,742) by randomly splitting the 297 NanoString nCounter cartridges into two groups. The training and validation cohorts are statistically indistinguishable from one another and from the overall trial cohort (**Supplementary Table 20**)^33^.

### RNA extraction

Five 4 µm formalin-fixed paraffin-embedded (FFPE) sections per case were deparaffinised, tumor areas were macro-dissected and RNA extracted according to Ambion® Recoverall™ Total Nucleic Acid Isolation Kit-RNA extraction protocol (Life Technologies™, Ontario, Canada) except that samples were incubated in protease for 3 hours instead of 15 minutes. RNA samples were eluted and quantified using a Nanodrop-8000 spectrophotometer (Delaware, USA). Samples, where necessary, underwent sodium-acetate/ethanol re-precipitation. We selected 33 genes of interest from key functional nodes in the PIK3CA signaling pathway^55^ and 6 reference genes (**Supplementary Table 16, Supplementary Results section 4**). Probes for each gene were designed and synthesized at NanoString® Technologies (Washington, USA). RNA samples (400 ng; 5 μL of 80 ng/μL) were hybridized, processed and analyzed using the NanoString® nCounter® Analysis System, according to NanoString® Technologies protocols.

### Univariate data analyses

In order to avoid dataset-specific bias, all included studies were analyzed independently (**Supplementary Table 2**). First, each dataset was pre-processed independently, as described in the ‘mRNA abundance data pre-processing’ section above. Next, genes across all the datasets were evaluated for their prognostic power using a univariate Cox proportional hazards model followed by the Wald-test (R package: survival v2.36-9). For breast, NSCLC and ovarian cancers with different survival end-points, overall survival (OS) was used as the survival time variable; for the studies that did not report OS, we used the closest alternative endpoint available in that study (*e.g*. disease-specific survival or distant metastasis-free survival). For colon cancer, all studies reported relapse/disease free survival and hence was used as survival end-point. All the genes were subsequently ranked by the Wald-test p-value within each study. The top genes across all studies were compared on multiple criterion (**Supplementary Results section 1**):

#### 1 Rank Product

The Rank Product^56^ of each gene was computed as:

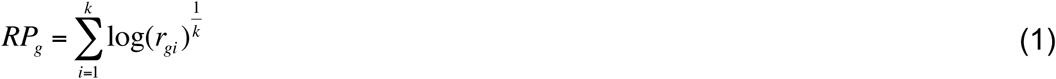

Here *k* represents the number of studies which had the mRNA abundance measure available for gene *g. r*_*i*_ is the rank of gene *g* in study *i*. The overall ranking table was used as a benchmark to identify datasets in which a given gene was ranked farthest when its rank product was compared to studywise ranks. The farthest dataset count was computed for the overall top ranked (100, 200, 300,…, 1000, 2000) genes (**Supplementary Figure 3a-e**).

#### 2 Percentile ranks

The p-value (Wald-test) based ranking was transformed into percentile ranks within each study. These ranks were used as a measure of gene’s position with reference to the benchmark rank derived in the step 1 to evaluate deviation of genes’ ranks for each study (**Supplementary Figure 3f-l**).

#### 3 Intra- and inter-study correlation

The mRNA abundance profiles of common genes across all studies were extracted and patient wise Spearman rank correlation coefficient was estimated (R package: stats v2.13.0). The correlation coefficient was used to further analyze intra- and inter-study correlation in order to identify any outlier studies (**Supplementary Figure 3j-l**).

### Eliminating redundant mRNA profiles (breast cancer data)

The Spearman rank correlation coefficient was also used to establish a non-redundant set of patients. This is important not only to identify any patients that might have participated in more than one study or duplicate data used in multiple papers, but also to train a robust model thereby preventing model over-fitting. The survival data of patients with high correlation coefficient (ρ ≥ 0.98) was matched, and we found 22 samples^57, 58^ having identical survival time and status. These patients were removed from further analyses (**Supplementary Figure 3m**).

### Meta-analysis

Following univariate analyses and elimination of redundant patients, the remaining studies were divided into two sets, training and validation (**Supplementary Tables 2-5**). The RMA normalized mRNA abundance measures were median scaled within the scope of each dataset (R package: stats v2.13.0).

#### 1 Gene hazard ratio

The hazard ratio for all the genes by combining samples from all the training datasets was estimated using the univariate Cox proportional hazards model. The Cox model was fit to the median dichotomized grouping of mRNA abundance profiles of the samples as opposed to continuous measure of mRNA abundance.

#### 2 Interaction hazard ratio

The hazard ratio for all the protein-protein interactions gathered from the NCI-Nature pathway interaction database were estimated using a multivariate Cox proportional hazards model. A Cox model, shown below, was fit to median dichotomized patient grouping of each of the interacting gene pairs:

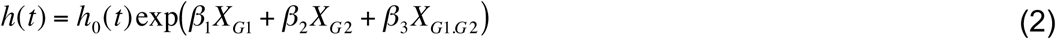

where X_G1_ and X_G2_ represent patient’s group for gene 1 and gene 2. X_G1.G2_ represents patient’s binary interaction measure between the gene 1 and gene 2, as shown below:

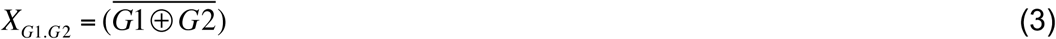

where ⊕ represents exclusive disjunction between the grouping of each gene. The expression encodes *XNOR* boolean function emulating *true (1)* whenever both the interacting genes belong to the same group.

### Subnetwork module-dysregulation score (MDS)

The pathway-based subnetworks were scored using three different models. These models compute a module-dysregulation score (MDS) by incorporating the hazard ratio of nodes and edges that form the subnetwork:

#### 1 Nodes + Edges

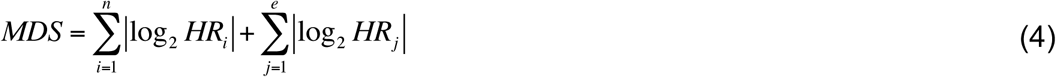

#### 2 Nodes only

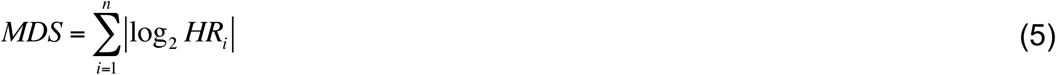

#### 3 Edges only

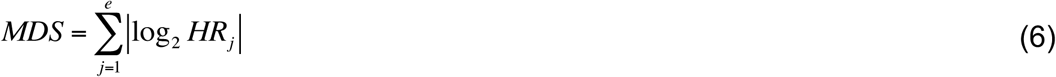

where *n* and *e* represent total number of nodes (genes) and edges (interactions) in a subnetwork respectively. *HR* represents the hazard ratios of genes and the protein-protein interactions in a subnetwork (P < 0.05) (section: Meta-analysis). The subnetworks were ranked according to their MDS, thereby identifying candidate prognostic features.

### Patient risk score

The subnetwork MDS was used to draw a list of the top *n* subnetwork features for each of the three models (section: Subnetwork module-dysregulation score). These features were subsequently used to estimate patient risk scores using Model N+E, N and E. The patient risk score for each of the subnetworks (*risk*_*SN*_) was expressed using the following models:

#### 1 Nodes + Edges

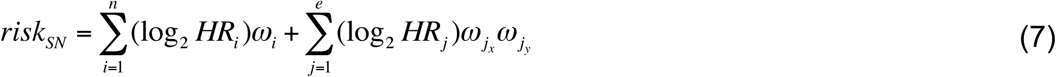

#### 2 Nodes only

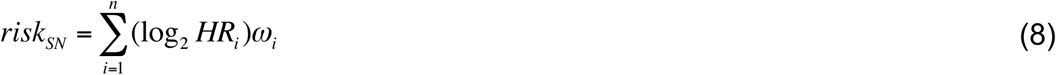

#### 3 Edges only

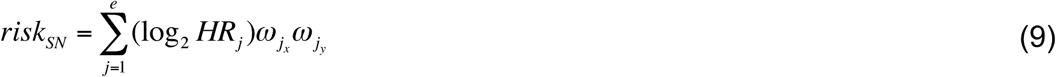

where *n* and *e* represent the total number of nodes (genes) and edges (interactions) in a subnetwork (*SN*), respectively. *HR* is the hazard ratio of genes and the protein-protein interactions (P < 0.5; only to filter out genes where Cox model fails to fit estimating large/unstable coefficients) (section: Meta-analysis) in a subnetwork. *x* and *y* are the two nodes connected by an edge *e*_*j*_ and ***ω*** is the scaled intensity of an arbitrary molecular profile (*e.g*. mRNA abundance, copy number aberrations, DNA methylation beta values etc). A univariate Cox proportional hazards model was fitted to the training set, and applied to the validation set for each of the subnetworks. The prognostic power of all three models was compared using non-parametric two sample Wilcoxon rank-sum test (R package: stats v2.13.0).

### Subnetwork feature selection

In order to narrow down the size of subnetwork features in each of the three models yet maintaining the prognostic power, we fitted a Cox model based Generalised Linear Model (*L1*-penalty) in 10-fold cross validation setting on the training cohort (R package: glmnet v1.9-8). SIMMS package supports additional machine learning algorithms including elastic-nets (ridge to lasso), backward elimination and forward selection (R package: MASS v7.3-12). The fitted coefficients (*β*) were subsequently used to estimate an overall measure of per patient risk score for the validation set using the following formula:

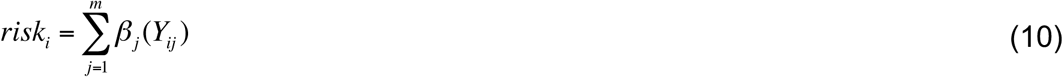

where *Y*_*ij*_ is the *i*^*th*^ patient’s risk score for subnetwork *j*. The training set HRs of the nodes and edges were used to compute *Y*_*ij*_ (section: Patient risk score). Next, we median dichotomized the validation cohort into low- and high-risk patients (or quartiles) using the median risk score (or quartiles) estimated on the training set. The risk group classification was assessed for potential association with patient survival data using Cox proportional hazards model and Kaplan-Meier survival analysis.

### Randomization of candidate subnetwork markers

Jackknifing was performed over the subnetwork marker space for four tumour types; breast, colon, NSCLC and ovarian. Ten million prognostic classifiers (200,000 for each size *n*=5,10,15,….,250; where *n* represents the number of subnetworks) were randomly sampled using all 500 subnetworks. The predictive performance of each random classifier was measured as the absolute value of the log_2_-transformed hazard ratio obtained by fitting a multivariate Cox proportional hazards model using Model N.

### Visualizations

All plots were created in the R statistical environment (v2.13.0). Forest plots were generated using rmeta package (v2.16), all others were created using the BPG (P’ng *et al*., in review), lattice (v0.19-28), latticeExtra (v0.6-16) and VennDiagram (v1.0.0) packages.

## List of abbreviations

AFFY: Affymetrix,
CI: Confidence interval,
DRFS: Distant metastasis free survival,
ER: Estrogen receptor 1,
FFPE: Formalin-fixed, paraffin-embedded,
Her2: human epidermal growth factor receptor 2,
HR: Hazard ratio,
IHC: Immunohistochemistry,
ILMN: Illumina,
MDS: module-dysregulation score,
NSCLC: Non-small cell lung cancer,
OS: Overall survival,
PgR: Progesterone receptor,
RP: Rank Product,
SIMMS: Subnetwork integration for multi-modal signatures,
SN: Subnetwork,
Tukey HSD: Tukey honestly significant difference,
TCGA: The cancer genome atlas.

## Author contributions

PCB and SH initiated the research project, and designed and implemented SIMMS. SH, MHWS, CQY, JW, FN and PCB collected and analyzed pan-cancer data. CQY and SH analyzed the NanoString data and performed the computational modeling and biomarker discovery in that data. MG performed subnetwork permutation analysis and contributed to the R package. NCM processed copy number data and provided statistical support. VSS, CD, CAC, CLB, CJHvdV, AH, DGK, CJM, LYD, CS, CWR and JMB initiated and designed all TEAM profiling experiments, and performed NanoString profiling on this cohort. SH, MHWS and PCB interpreted pan-cancer results; SH, JMB and PCB interpreted TEAM results. AK and PL provided intellectual input in algorithm development. SH and PCB wrote the manuscript with contributions from all authors.

## Acknowledgements

We gratefully acknowledge the support of all pathologists, treating physicians and the participation of all patients who consented to provide paraffin blocks for the study. We thank Nicole Carpe, Kristen Geras, Dr. Jane Starczynski, Mary Anne Quintayo, Bei Jiang and Sally Stasi for technical assistance (OICR) and Jacqueline Stephen and Tammy Piper (University of Edinburgh) for database assistance. The authors thank all members of the Boutros lab for insightful suggestions, in particular Christine P’ng for data visualization and Daryl Waggott for bioinformatics support in data normalization. The authors thank Drs. Lincoln D. Stein (OICR), Gary D. Bader (University of Toronto), Guanming Wu (OICR) for insightful comments.

This study was conducted with the support of the Ontario Institute for Cancer Research to AK, JMB and PCB through funding provided by the Government of Ontario, and with the support of the Canadian Breast Cancer Foundation (CBCF) to CQY. PCB was supported by a Terry Fox Research Institute New Investigator Award and a CIHR New Investigator Award. This study makes use of data generated by the Molecular Taxonomy of Breast Cancer International Consortium, which was funded by Cancer Research UK and the British Columbia Cancer Agency Branch. The results published here are in whole or part based upon data generated by The Cancer Genome Atlas pilot project established by the NCI and NHGRI. Information about TCGA and the investigators and institutions who constitute the TCGA research network can be found at http://cancergenome.nih.gov/.

